# Introgression obscures and reveals historical relationships among the American live oaks

**DOI:** 10.1101/016238

**Authors:** Deren A. R. Eaton, Andrew L. Hipp, Antonio González-Rodríguez, Jeannine Cavender-Bares

## Abstract

Introgressive hybridization challenges the concepts we use to define species and infer phylogenetic relationships. Methods for inferring historical introgression from the genomes of extant species are now widely used, however, few guidelines have been articulated for how best to interpret results. Because these tests are inherently comparative, they are sensitive to the effects of missing data (unsampled species) and non-independence (hierarchical relationships among species). We demonstrate this using genomic RADseq data sampled from all extant species in the American live oaks (*Quercus series Virentes*), a group notorious for hybridization. By considering all species, and their phylogenetic relationships, we were able to distinguish true hybridizing lineages from those that falsely appear admixed. Six of seven species show evidence of admixture, often with multiple other species, but which is explained by hybrid introgression among few related lineages occurring in close proximity. We identify the Cuban oak as the most admixed lineage and test alternative scenarios for its origin. The live oaks form a continuous ring-like distribution around the Gulf of Mexico, connected in Cuba, across which they could effectively exchange alleles. However, introgression appears highly localized, suggesting that oak species boundaries, and their geographic ranges have remained relatively stable over evolutionary time.

## Introduction

Introgressive hybridization is a common phenomenon among biological organisms, including our own species (Green *et al*. 2010). It impacts how we understand the nature of species and infer their historical relationships, with important implications for conservation and biodiversity research (Rhymer & Simberloff 1996). Because introgression between divergent lineages can give rise to genetically admixed individuals and populations that are heterogeneously distributed in space and/or time (Avise 2000, Petit & Excoffier 2009), sampling such individuals will generally bias estimates for the order and timing of species divergences (Leaché *et al.*. 2014). Yet phylogenetic studies rarely sample a sufficient number and variety of individuals to detect whether admixture is present, or variable within species. Similarly, the common practice of excluding apparent hybrid individuals from phylogenetic studies prevents researchers from evaluating their influence on phylogeny. To the extent that introgression is common, the practice of sparse sampling in phylogenetics will underestimate its frequency, and in doing so infer an inflated role for stochastic processes, such as incomplete lineage sorting (Maddison & Knowles 2006), in explaining discordant genealogical relationships.

Recent years have seen the development of new methods for inferring admixture from the genomes of extant species (Green *et al*. 2010, Durand *et al*. 2011), the results from which are often interpreted as evidence of hybrid introgression between their ancestors. Connecting pattern (admixture) and process (introgression) in this way is a difficult problem, however, and one that similarly suffers from the effects of sparse taxon sampling. To account for such effects, we highlight two important considerations that should generally be taken into account. First, the problem of missing samples: when the true source of introgression is not sampled (i.e., it is a ghost lineage) the source will usually be incorrectly attributed to the sampled population most closely related to the ghost lineage (Durand *et al*. 2011, Eaton & Ree 2013, Rogers & Bohlender In Press). In practice, the extent to which truly spurious conclusions would be drawn from sampling a closest available (or extant) lineage will generally depend on the size of the clade to which hybridizing lineages belong, and their rate of ecological or morphological divergence. Diverse clades would require very dense sampling to identify that a species or population that appears admixed does not have a close relative harboring a yet stronger signal of admixture.

A second and related consideration is that even when all relevant lineages are sampled in a study, it still remains difficult to distinguish a history of introgression between two populations from a signal of admixture between those populations that can arise when one species harbors introgressed alleles from a close relative of the other (Eaton & Ree 2013). To distinguish true introgression from such secondary genomic admixture, introgression must be considered in an explicitly hierarchical (phylogenetic) context, rather than on a species-by-species basis. For example, suppose there are two species, A and D, which exchanged alleles at some time in the past. Species A is member of a clade including several other species (B and C) with which it shares many derived alleles since their divergence from D. As a consequence of their relatedness, introgression from species A into D will necessarily introduce alleles that it also shares with its close relatives, which can give the appearance (admixture) that B and C also hybridized with D. To identify whether the relatives of A independently introgressed into D, versus whether they simply share ancestry with the true hybridizing lineage, requires not only sampling all relevant lineages in the clade, but also accounting for their phylogenetic structure.

Oaks (*Quercus*) are notorious for hybridization (Hardin 1975, Burger 1975) to the extent they have been dubbed a “worst case scenario for the biological species concept” (Coyne & Orr 2004). For this reason, they also provide a compelling case study for investigating introgression at the clade level, among multiple interacting species. Within the genus, the American live oaks (*Quercus* section Virentes Nixon) form a young clade of seven ecologically divergent species that span a range of climatic regimes from the seasonal dry tropics to the temperate zone (Muller 1961, Nixon 1984, Cavender-Bares *et al*. 2011; In Press). They include both narrow endemics and widespread species that collectively cover the southeastern US, eastern Mexico, southern Baja, Central America, and Cuba (Fig. 1A). The species are all diploid and interfertile, and many occur in sympatry throughout all or parts of their range. A complex history of hybridization has likely contributed to difficulties in resolving their phylogenetic relationships (Cavender-Bares & Pahlich 2009, Gugger & Cavender-Bares 2013).

**Figure 1:**
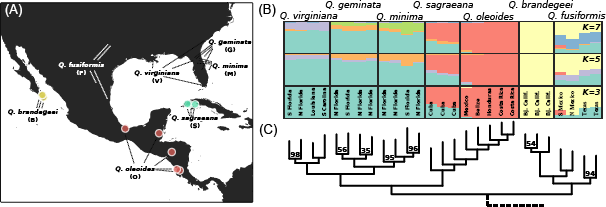
Sampling locations spanning the geographic ranges of each of the seven live oak taxa. (B) Population clustering inferred with admixture at three values of *K*. Sampling locations are indicated. (C) Rooted ML phylogeny inferred from the largest (Allmin4) concatenated RADseq data set. Only ingroup taxa are shown. Bootstrap support is 100 except where indicated.

The live oaks are part of a predominately American oak clade (Hipp *et al*. 2014, Pearse & Hipp 2009) comprising sections *Quercus* (the white oaks *sensu stricto*, including the live oaks of the Americas and roburoids of Eurasia), *Lobatae* Loudon (the red or black oaks), and *Protobalanus* (Trelease) A.Camus (the intermediate or golden oaks). The red and white oak clades became morphologically distinct ca. 23–33 (Borgardt & Pigg 1999). Although hybrids are commonly observed within each major section (Hardin 1975), hybrid swarms are uncommon, as is hybridization between major sections (Muller 1961). The live oaks are sister to the remainder of the white oaks, making them phylogenetically distant and isolated from all other oak species, and thus a manageable system in which to reconstruct a clade-level history of introgression.

Here we utilize restriction-site associated DNA sequencing (RADseq) (Baird *et al*. 2008) to sample thousands of genomic regions across a large number of samples for phylogenetic inference, and to test introgression between lineages. A recent study demonstrating high conservation of RAD sequences across a phylogenetic scale spanning more than 40 Mya in the American clade oaks (Hipp *et al*. 2014) motivates our current study. While genetic admixture has been previously described in the live oaks between focal species pairs (Cavender-Bares & Pahlich 2009, Gugger & Cavender-Bares 2013), this is the first study to bring genome-scale data to bear on the question, and more importantly, to investigate introgression among all extant species in the clade simultaneously and within a phylogenetic context.

We focus particular attention to resolving the phylogenetic placement of the Cuban oak species, *Q. sagraeana*. The origin of this isolated and distinct taxon has long puzzled systematists: its origin has been variously ascribed to one or more species in Florida, to a Central American species, or to hybridization among other live oaks (Muller 1961, Nixon 1984, Gugger & Cavender-Bares 2013). Chloroplasts are commonly exchanged between sympatric oak species (Whittemore & Schaal 1991, Petit *et al*. 1997), and consequently chloroplast DNA (cpDNA) haplotypes exhibit little species specificity compared to nuclear markers (Petit & Excoffier 2009, Dumolin-Lapegue *et al*. 1999). The cpDNA haplotype common in Cuba is also shared with both of its hypothesized parent lineages, and is thus inconclusive about the biogeographic origins of the species (Gugger & Cavender-Bares 2013). Using >70K RAD loci sequenced from multiple individuals across the geographic ranges of all seven extant species of live oaks, we ask the following: (1) Which lineages have experienced hybrid introgression? (2) How does admixture affect phylogenetic inference? (3) Can we tease apart non-independent signals of admixture among multiple closely related species? And (4) what is the origin of the Cuban oak?

## Materials and Methods

### Sampling

Four to five individuals were sampled from across the geographic range of each of the seven live oak species for RAD sequencing (Fig. 1A), in addition to seven outgroup samples (Four non-*Virentes* white oaks: *Q. engelmannii, Q. arizonica, Q. durata, Q. douglasii*; one golden oak: *Q. chrysolepis;* and two red oaks: *Q. nigra, Q. hemisphaerica*). Leaf samples were collected from wild plants (live oaks) or plants grown in the University of Minnesota greenhouse (outgroup samples). Identification to species was based on leaf, bark, and stem height characters following Muller (1961), Kurz & Godfrey (1962), and Nixon & Muller (1997). Leaves were collected from wild plants in the field, maintained fresh during transport, and stored at -80C until extraction. Voucher specimens for all RAD sequenced individuals are housed in the University of Minnesota Bell Museum of Natural History (Table S1).

### RADseq preparation and sequencing

DNA was extracted from fresh or frozen material using the DNeasy plant extraction protocol (DNeasy, Qiagen, Valencia, CA) as reported in Cavender-Bares & Pahlich (2009). RAD libraries were prepared by Floragenex Inc. (Eugene, Oregon) using the PstI restriction enzyme and sonication following the methods of Baird *et al*. (2008). An initial multiplex library was created from 30 barcoded and pooled samples sequenced on an Illumina GAIIx sequencer to generate 100 bp single end reads. To increase coverage a second library was prepared that included an additional 15 samples, seven of which were technical replicates of samples in the first library, sequenced on an Illumina HiSeq 2000 to generate 100 bp single end reads. After an initial analysis to check that technical replicates grouped together in phylogenetic analyses, they were combined, except for one replicate that may have been contaminated and was excluded. Two additional samples were discarded during bioinformatic analyses due to low sequencing coverage (“TXVW2” and “CUMM5”) resulting in 34 final samples.

### RAD*seq assembly*

Data were assembled into *de novo* loci using *py*RAD v.2.13 (Eaton 2014). Quality filtering converted base calls with a score <20 into Ns and reads with >5 Ns were discarded. Illumina adapters and fragmented sequences were removed using the filter setting “1” in *py*RAD. Filtered reads were clustered at two different thresholds for within-sample clustering, 85% and 92%, both of which yielded similar results, therefore we report only the 85% run. Error rate and heterozygosity were jointly estimated from aligned clusters for each sampled individual and the average parameter values were used when making consensus base calls. Clusters with a minimum depth of coverage <5 were excluded. Loci containing more than two alleles after error correction were excluded as potential paralogs (all taxa in this study are diploid). Consensus loci were then clustered across samples at 85% similarity and aligned. A final filtering step excluded any loci containing one or more sites that appear heterozygous across more than five samples, as we suspect this is more likely to represent a fixed difference among clustered paralogs than a true polymorphism at the scale of this study. The final assembly statistics appeared robust to the choice of filtering thresholds.

In addition to assembling full data sets, smaller matrices were also assembled in which taxa from one or two major clades were selectively excluded. This allowed phylogenetic inference to be performed separately for each major clade in the live oaks, rooted by the outgroups, but without the influence of shared SNPs between taxa from distant ingroup clades. The motivation for this approach is that to the extent introgression has introduced synapomorphies between distant relatives, subsampling will censor their effect, making them appear instead as autapomorphies (Eaton & Ree 2013). To explore the effect of missing data we also assembled each data set with different minimums for sample coverage (the number of samples for which data must be recovered to include a RAD locus in the data set). A large but incomplete version required at least four samples have data for a locus (e.g., “Allmin4”), while a smaller more complete version was also assembled (e.g., “Allmin20”). In total, 15 data sets were generated. The source of missing data between samples was investigated using Mantel tests (9999 permutations) that measured the Spearman's rank correlation between the Jaccard's distance of the proportion of shared loci between samples, pair-wise phylogenetic distance, and number of raw input reads.

### Phylogeny and population clustering

For each assembled data set RAD loci were concatenated and missing data entered as Ns to create a phylogenetic supermatrix. Maximum likelihood (ML) trees were inferred in RAxML v.7.2.8 (Stamatakis 2014) with bootstrap support estimated from 200 replicate searches from random starting trees using the GTR+Г nucleotide substitution model.

To better visualize genomic variation within individuals we inferred population clustering with admixture from SNP frequency data within the program *Structure* v.2.3.1 (Pritchard *et al*. 2000). To minimize missing data across individuals we used 14,011 putatively unlinked bi-allelic SNPs, sampled by selecting a single SNP from each locus in the “Ingroupmin20” data set (17% missing data), which includes only ingroup samples and requires that a locus contain data for at least 20 samples. Ten replicates were run at each value of *K* between 2-8. Each run had a burn-in of 50K generations followed by 500K generations of sampling. Replicates were permuted in the program *CLUMPP* (Jakobsson & Rosenberg 2007), and the optimal *K* was inferred using the online resource *StructureHarvester* (Earl & vonHoldt 2012).

We also used the program *TreeMix* (v.1.12; Pickrell & Pritchard 2012) to jointly estimate a tree topology (or graph) with admixture using pooled SNP frequency data. For this, individuals were pooled into populations matching to species designations except for *Q. fusiformis* which was split into separate populations for samples from Mexico and Texas. The four non-*Virentes* white oak samples were pooled as an outgroup population. A single bi-allelic SNP was randomly sampled from each variable locus that contained data for at least one individual across all populations, yielding a total of 12,061 bi-allelic SNPs. We inferred a topology without admixture, as well as when allowing between 1-5 admixture events.

### Introgression analyses

The four-taxon D-statistic (Durand *et al*. 2011) is a well-known metric for detecting admixture between diverged lineages based on the frequencies of SNPs that are discordant with a hypothesized species tree topology. It was most notably used to demonstrate introgression between Neanderthals and modern humans from full genome data (Green *et al*. 2010), and has similarly been applied to non-model organisms using RADseq data (The Heliconius Genome Consortium 2012, Eaton & Ree 2013). Given a four-taxon pectinate tree [(((P1,P2),P3),O)] in which the outgroup/ancestral allele is labeled “A”, and a derived allele labeled “B”, the D-statistic compares the occurrence of two discordant site patterns, ABBA and BABA, representing sites in which an allele is derived in P3 relative to O, and is derived in one but not both of the sister lineages P1 and P2. These discordant sites can arise through the sorting of ancestral polymorphisms, but will generally do so with equal frequency due to the stochastic nature of this process. Alternatively, they may arise if introgression occurs between P3 and either P2 or P1, in which case one site pattern will occur more frequently than the other. The D-statistic provides a test for historical admixture by calculating asymmetry in the relative occurrence of these two discordant site patterns:

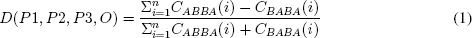

where *C*_*ABBA*_(i) and *C*_*BABA*_(i) are indicator variables of 0 or 1 depending on whether ABBA or BABA is present at each site. Following Durand *et al.* (2011), we used SNP frequencies instead of allele counts in this study to allow for the inclusion of heterozygous sites. Thus, *D* was calculated as:

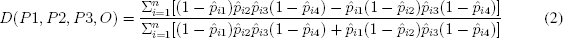

where 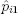 is the frequency of the derived allele in taxon P1 at site *i.* If the sampled individual has both copies of the derived allele at this site 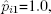 if it is heterozygous 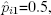 otherwise 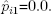 We calculated *D* over all combinations of four taxa fitting the maximum likelihood topology as well as alternative topologies of interest. For ingroup taxa we iterated over each sampled individual separately, but for the outgroup taxon instead used a pooled group of samples to measure the SNP frequency. This was made up of the four non-*Virentes* white oak samples, with 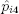 calculated as the frequency of derived alleles in all 2N locus copies for N outgroup individuals containing data for a given site. This allowed us to maximize the use of RADseq data with missing sequences, since we could use any locus for which the three sampled ingroup taxa shared data with at least one outgroup. This approach also has the effect of down-weighting D if the ancestral allele is not fixed across multiple outgroup samples, making it a more conservative test.

For each test we measured the standard deviation of *D* from 200 bootstrap replicates in which RAD loci were re-sampled with replacement to the same number as in the original data set, as in Eaton & Ree (2013). The observed *D* was converted to a Z-score measuring the number of standard deviations it deviates from 0, and significance was assessed from a P-value using α=0.01 as a conservative cut-off after Holm-Bonferoni correction for multiple testing (number of possible sample combinations fitting the given species tree hypothesis).

Partitioned D-statistics (Eaton & Ree 2013) are an extension to this test relevant at deeper evolutionary time scales where the P3 lineage may include multiple distinct sub-lineages with independent histories of introgression. It measures a five-part allele pattern [(((P1,P2),(P3_1_,P3_2_)),O)], and contrasts two P3 sub-lineages at a time by measuring *D* for three separate pairs of allele counts (ABBBA/BABBA, ABBAA/BABAA, and ABABA/BAABA). These statistics measure asymmetry in the occurrence of derived alleles present in both P3 sub-lineages (D_12_), only P3_1_ (D_1_), or only P3_2_ (D_2_), and present in P2 or P1 but not both (Fig. 2A).

**Figure 2:**
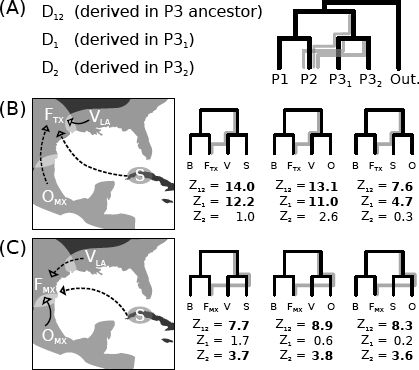
Teasing apart non-independent signals of admixture. (A) Partitioned D-statistics test for directional introgression from the P3 lineage into P2 or P1 and contrast P3 sub-lineages as introgressive donors. Results are reported as Z-scores. (B) Three closely related lineages (S, V & O; taxon names abbreviated as in Fig. 1) each share alleles with F in Texas to the exclusion of B (significant *D*_12_, but when contrasted against each other (D_1_ and D_2_) only V shares uniquely introgressed alleles with F_*T X*_ relative to the other two P3 sub-lineages. (C) A similar test examining F from coastal Mexico shows the opposite result: F_*MX*_ only shares uniquely introgressed alleles with O, while apparent admixture between F_*MX*_ and S or V is a consequence of the shared ancestry of O with S and V.

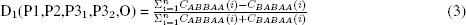

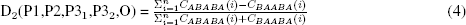

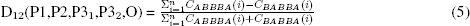

As in the four-taxon tests, we used the four non-*Virentes* white oak samples to represent the outgroup, and used a SNP frequency-based version of the test to include data for heterozygous individuals. All D-statistics were measured in pyRAD v.2.13.

In contrast to the four-taxon D-statistic, the partitioned test is polarized by defining P3 as a donor lineage, and P2 or P1 as recipients, which allows D_12_ to act as an indicator of the direction of introgression. Briefly, consider a case where introgression occurred in the reverse direction from how we assign samples to the tips of the tree (e.g., from P2 into P3_1_); in this case, P3_2_ would not contain the same derived alleles that P2 shares with P3_1_ through introgression, and thus the indicator variable D_12_ would be non-significant, indicating introgression did not occur in this direction. If we then swap samples across the tips to re-define the P3 lineage, such that introgression occurred from the defined P3_1_ sub-lineage into P2, we would now find that P3_2_ also shares many of the same introgressed alleles that P3_1_ shares with P2 (significant D_12_), due to the fact that many of these alleles arose in the ancestor of the two sampled P3 sub-lineages. In addition to indicating directionality, partitioning ancestral alleles from those that are derived uniquely to either P3 sub-lineages also allows us to distinguish whether introgression occurred from each P3 sub-lineage independently into P1 or P2, or if it occurred from only one (Eaton & Ree 2013). We apply this test to two separate cases in the live oaks, involving *Q. fusiformis* and *Q. sagraeana*, in which four-taxon tests show evidence of admixture involving more than two taxa, to test whether each taxon pair hybridized independently.

### Demographic models

To investigate the origin of the Cuban oak we compared the joint site frequency spectrum (SFS) generated under three demographic isolation-migration models (Fig. 4A) to that in our observed data, with a focus on SNPs segregating within and between populations of *Q. oleoides, Q. sagraeana,* and the Florida oaks clade, using the program ∂a∂i (Gutenkunst *et al*. 2009). Data were pooled for the three closely related species in Florida, and the SFS was projected down to require that every locus contain data for at least five individuals in Florida, three individuals in *Q. oleoides*, and three individuals in *Q. sagraeana* (projected chromosomes = [10,6,6]). A single bi-allelic SNP was randomly selected from each variable locus, yielding 1,626 SNPs from 7,794 usable loci after data projection.

**Figure 4:**
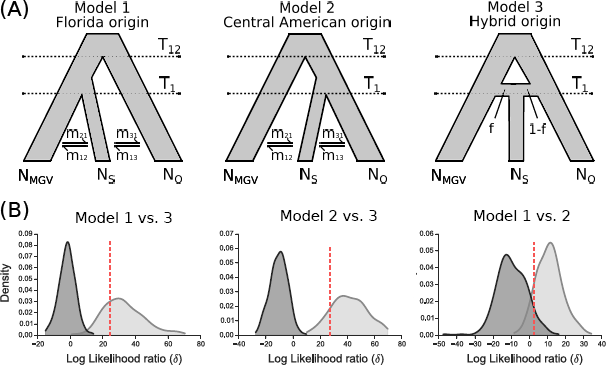
Three demographic models for the origin of the Cuban oak (S). (A) In models 1 and 2 (9 parameters) S is derived from one mainland taxon or the other (O or MGV; taxon names are abbreviated as in Fig. 1) with subsequent migration between Cuba and either mainland lineage. In model 3 (7 parameters) S forms through instantaneous admixture (hybrid speciation) and remains isolated thereafter. (B) Results of Monte Carlo model comparisons. Distributions of likelihood ratios (*δ*) show the difference in fit between models when data are simulated under one model or the other. The likelihood ratio fit between models for our observed data is shown in red (*δ*_*obs*_). The proportion of the null model's *δ* distribution (dark grey) to the right of *δ*_*obs*_ measures the false positive rate, and the proportion of the alternative model's *δ* distribution (light grey) that overlaps with the null distribution measures the power to reject the null. Model 2 is the best fit to our observed data.

The first two demographic models have 9 parameters and differ only in their topology: in model 1 the Cuban oak is derived from Florida, while in model 2 it originates from Central America (Fig. 4A). Model parameters include an effective population size for each population (*N*_*MGV*_, *N*_*O*_, and *N*_*S*_) and migration rates between adjacent populations (*m*_12_, *m*_21_, *m*_23_, *m*_32_). At time *T*_*2*_, two ancestral populations diverge (viewed forward in time), and at time *T*_*1*_ the Cuban population diverges from its sister lineage to maintain a separate constant population size. Model 3 has only 7 parameters. In this model, *T*_*2*_ is again the divergence time for two ancestral populations, but *T*_*1*_ is now an event in which an independent Cuban population is formed by an instantaneous fusion of a proportion (*f*) of the Florida population and (1-*f*) of *Q. oleoides*. There is no further migration between populations.

We used the log L-BFGS-B optimization method to fit parameters for each model. Searches were started from 10 randomly perturbed starting positions, for a maximum of 5 iterations, followed by a final search using the best-inferred parameters from the previous step as a starting position for a maximum of 20 additional iterations. Extrapolation was performed with a grid size of [12,20,32]. To attain confidence intervals on parameter estimates we performed parametric bootstrapping by simulating 200 data sets for each of the three models using the program *ms* (Hudson 2002). Bootstrap SFS data were simulated under their ML estimated parameter values and then re-optimized in ∂a∂i to estimate the parameters that would generate these data under the same model by which they were generated.

The same simulated data sets were also used for Monte Carlo model selection (Boettiger *et al*. 2012). Here, in addition to fitting the simulated data sets to the model under which they were simulated, each data set was also fit to the other two models (9 model fits total), and for each comparison a likelihood ratio [*δ* = -2(log L_0_ - log L_1_)] was calculated. Larger values for *δ* indicate more support for model 1 relative to model 0(the null). Our goal in model selection is to calculate how big *δ* should be in order to decide that model 1 is closer to the truth than model 0 (Boettiger *et al*. 2012). Power to distinguish models, and the sensitivity of our tests, were assessed from the overlap in distributions of *δ* values from simulated data, and their comparison to δ for our observed data.

### Reproducibility

Scripts to download archived sequence data (NCBI: PRJNA277574), assemble it, and reproduce all analyses in this study are compiled into IPython notebooks (Pérez & Granger 2007), a tool for reproducible science, available at https://github.com/dereneaton/virentes (doi:xxxyyy).

## Results

### RAD data assembly

Following quality filtering and clustering (85% similarity) 77M raw reads (mean±S.D. 2.13M±1.75M per sample) were reduced to an average of 57K±25K high coverage stacks per sample, with a mean depth of 23X. These were further filtered to 52K±22K consensus sequences per sample (Table S1). Data sets that were assembled with different minimums for sample coverage or with samples excluded had different proportions of missing data: The largest but most incomplete assembled data matrix that includes all loci shared across at least four samples (Allmin4) has 55.5% missing data for 34 individuals across 78,727 loci, while all other matrices have fewer missing data (9.6–52.1%; Table 1).

**Table 1:**
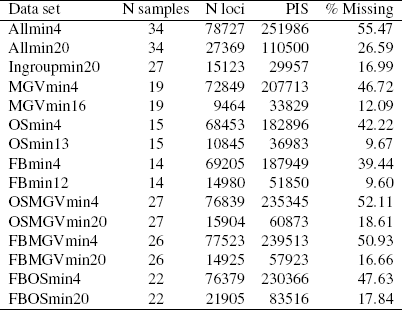
Size, completeness, and the number of phylogenetic informative sites (PIS) in 15 assembled RADseq data sets.

The distribution of missing data did not show strong hierarchical structure, as would be expected if most missing data was caused by locus dropout due to the disruption of restriction recognition sites (Fig. S1). Instead, for the largest data set (“Allmin4”) the mean number of raw reads was a better predictor for the number of shared loci between samples than was the phylogenetic distance between samples (Mantel r_ρ_=0.372, *P*=0.010, and r_ρ_=-0.145, *P*=0.240, respectively). A similar result was observed in the more complete “Allmin20” data set (Mantel r_ρ_=0.479, *P*=0.002, and r_ρ_=0.087, *P*=0.523, respectively), suggesting that sequencing effort had a more significant impact on missing data than relatedness.

### Phylogeny

Missing data (the sparseness of concatenated matrices) had little effect on phylogenetic inference as the larger and more incomplete versions of each data set yielded similar or identical topologies to the smaller more complete version of that matrix (e.g., Allmin4 & Allmin20; Fig. S2), the latter often with lower bootstrap supports. All phylogenetic analyses recovered perfect support for three major clades: a Florida clade (*Q. minima*, *Q. geminata*, and *Q. virginiana*), a southwestern clade (*Q. brandegeei* and *Q. fusiformis*), and a Central American clade (*Q. oleoides* and *Q. sagraeana)* (Fig. 1C). Selectively excluding taxa sometimes yielded different relationships within each major clade, as expected if synapomorphies that are derived from introgression between lineages affect phylogenetic inference (Eaton & Ree 2013). For example, *Q. fusiformis* appears paraphyletic with respect to its putative sister taxon *Q. brandegeei* in data sets that include samples from all three major clades, but monophyly of *Q. fusiformis* is supported when the two other live oak subclades are excluded (Fig. S2E). A similar pattern is observed for the three Florida clade oaks, where Q. virginiana appears sister to the other two species in full data sets, but *Q. minima* is sister to the other two species when the southwest and Central American clades are excluded (Fig. S2G). The phylogenetic instability of *Q. virginiana* and *Q. fusiformis* is consistent with further evidence below that they have exchanged genes in Texas where they occur in sympatry and that this affects their phylogenetic placement.

### Population structure

Population clustering analyses revealed substantial heterogeneity in proportions of admixed ancestry within and between species. The best supported model (*K* =3) clustered populations into the same three major clades described above. The three oak species of the Florida clade are indistinguishable at low values of K (the number of distinct clusters) (Figs. 1B & S3), and much of their common ancestry is also shared through apparent admixture with both of their geographically adjacent taxa: *Q. fusiformis* in Texas to the west and *Q. sagraeana* in Cuba to the south. *Quercus sagraeana* also shares significant ancestry with *Q. oleoides* from Central America. In the southwest, *Q. fusiformis* shares ancestry with *Q. brandegeei* and *Q. virginiana*. In contrast, *Q. oleoides* forms a nearly distinct cluster, except for the sample from Mexico which shows slight admixture with different groups at different K values. Only *Q. brandegeei*, endemic to southern Baja California, forms a distinct non-admixed cluster in all analyses above K=2, suggesting it has remained genetically isolated from all other populations sampled in our study. Within each species, individuals with the greatest proportions of admixed ancestry appear as the earliest diverging in their clade (Fig. 1B-C), suggesting that inferred population-level relationships may reflect admixture proportions to a greater degree than they do historical population divergences - a major concern for phylogeographic studies below the species level.

### Treemix

*TreeMix* recovered the same topology for population-level relationships as our concatenated ML analyses performed on individuals. With the addition of one admixture edge, approximately 40% admixed ancestry is inferred between *Q. sagraeana* and a Florida clade oaks lineage, which also changes the backbone topology such that *Q. oleoides* is supported as sister to the remaining live oaks (Fig. S4). Adding a second admixture edge returns a graph similar to that of the original tree topology, but with admixture between *Q. virginiana* and *Q. sagraeana* (47% ancestry), and between *Q. virginiana* and *Q. fusiformis* in Texas (24% ancestry). A notable result of the latter edge is its effect on *Q. brandegeei*, which becomes no longer nested within *Q. fusiformis*. This shows how, despite being completely isolated from admixture itself, introgression occurring into a close relative of *Q. brandegeei* can still affect its phylogenetic placement.

The first admixture edge increases the log-likelihood (LL) by 68.2, the second edge by 60.6, while a third edge increases the LL by only 12.2, and all additional edges by less than 5. The first two inferred edges are concordant with D-statistic results reported below, and support admixture between *Q. virginiana* and both Q. fusiformis in Texas and *Q. sagraeana* in Cuba. The third inferred edge (Fig. S4), which shows admixture between the outgroup population and *Q. minima*, provides only a small improvement to the LL score and is not strongly supported by D-statistic results.

### D-statistics

Non-parametric D-statistics (ABBA-BABA tests) revealed substantial heterogeneity in the presence of admixture within and between species (Table 2). Few tests detected admixture uniformly across all iterations of sampled individuals. Significant results were largely limited to samples that occurred in close geographic proximity. For example, among the three sympatric oaks species in Florida, *Q. virginiana* shares derived alleles with *Q. geminata* to the exclusion of *Q. minima* when *Q. minima* is sampled from southern Florida, but not when sampled from northern Florida; an apparent consequence of all three taxa being more homogenized in the north (tests 1-5, Table 2). *Q. virginiana* is the only species in this clade to occur widely outside of Florida; however, it shows the same genetic similarity to the other two species in sympatry as it does in allopatry (tests 6 & 7, Table 2), suggesting that *Q. virginiana* has not received introgression from either species in the very recent past. Under an alternative topology in which Q. minima is sister to the other two Florida clade live oak species, we detect negligible admixture between *Q. virginiana* and *Q. geminata*, but admixture of both with the more rare taxon Q. minima (tests 1-4 & 6-8, Table 2). The most admixed sample of *Q. minima* groups with *Q. geminata* in several phylogenetic analyses (Fig. S2). Both Q. geminata and *Q. virginiana* are admixed with *Q. sagraeana* in Cuba, and *Q. virginiana* is also admixed with Q. fusiformis in Texas (tests 16 & 18-22, Table 2). Despite this, the three live oak species in Florida show little genetic differentiation from each other, and thus for simplicity we refer to them as a single pooled taxon (called the Florida clade, or abbreviated MGV) in several further analyses.

**Table 2:** Four-taxon D-statistic tests for admixture. Taxon names are abbreviated as in Fig. 1 and arranged such that ABBA>BABA. Outgroups not shown.

| Test | P1 | P2 | P3 | range $Z^a$ | nSig/N<sup>b</sup> |
| --- | --- | --- | --- | --- | --- |
| 1 | G | G | M | (0.0, 2.3) | 0/23 |
| 2 | M | M | G | (1.3, **6.8**) | 12/23 |
| 3 | G | G | V | (0.2, 2.4) | 0/17 |
| 4 | M | M | V | (0.2, **4.7**) | 7/17 |
| 5 | M | G | V | (0.1, **7.9**) | 28/47 |
| 6 | V | V | M | (0.0, 1.6) | 0/11 |
| 7 | V | V | G | (0.1, 2.5) | 0/11 |
| 8 | V | G | M | (0.0, **3.9**) | 1/11 |
| 9 | O | S | (MGV) | (**3.1, 16.2**) | 164/164 |
| 10 | (MGV) | S | O | (**14.7, 36.4**) | 164/164 |
| 11 | (MGV) | O | S | (**6.8, 25.8**) | 164/164 |
| 12 | O | O | F | (0.0, 1.6) | 0/39 |
| 13 | O | O | B | (0.1, 2.4) | 0/29 |
| 14 | B | F | O | (0.0, **8.1**) | 29/59 |
| 15 | B | F | S | (0.9, **8.1**) | 30/35 |
| 16 | B | F | (MGV) | (1.3, **17.9**) | 119/131 |
| 17 | B | B | F | (0.2, 2.6) | 0/11 |
| 18 | S | S | (MGV) | (0.0, **4.1**) | 2/32 |
| 19 | M | V | S | (1.1, **7.1**) | 17/35 |
| 20 | M | G | S | (0.0, **6.9**) | 18/47 |
| 21 | V | G | S | (0.0, 2.9) | 0/35 |
| 22 | (MG) | V | $F_{TX}$ | (**3.6, 10.6**) | 47/47 |
| 23 | O | O | $F_{MX}$ | (0.0, 1.7) | 0/19 |
| 24 | S | S | O | (0.0, **4.1**) | 2/14 |
| 25 | O | O | S | (0.1, **7.3**) | 10/29 |

<sup>a</sup> Bold indicates significance at $\alpha=0.01$ .
<sup>b</sup> Significant tests over possible sampled individuals.

The Cuban oak, *Q. sagraeana*, shows clear admixture with one or more Florida clade species and with *Q. oleoides* in Central America. Of the three possible rooted topologies for these three lineages (tests 9-11, Table 2) admixture is greatest when *Q. sagraeana* is sister to the Florida oaks clade (in conflict with our phylogenetic results) and exchanging genes with *Q. oleoides*. Here we see that *Q. sagraeana* shares more derived alleles, to the exclusion of the Florida clade, with the southernmost populations of *Q. oleoides* (Costa Rica & Honduras) than with northern populations (Mexico & Belize). The alternative test that is concordant with our phylogenetic results entails less admixture, meaning that *Q. sagraeana* shares more alleles with *Q. oleoides* than it does with the Florida clade oaks. We suspect that the third possible topology, in which Q. sagraeana diverged first from the other two species is unlikely, since *Q. sagraeana* exhibits little independent ancestry relative to the other two lineages (Fig. 1B).

*Quercus fusiformis*, which ranges from northern Mexico to eastern Texas, shows evidence of admixture with both of the other two major live oak clades, thus spanning the deepest splits in the tree. In Mexico it occurs in sympatry with *Q. oleoides*, and the two form a clear morphological hybrid zone (Cavender-Bares et al. In Press). We did not directly sample this hybrid zone in our genomic data set, however, the most geographically proximate samples from each taxon show evidence of admixture, suggesting introgression from *Q. oleoides* into *Q. fusiformis* (tests 12-14 & 23, Table 2). In Texas the range of *Q. fusiformis* overlaps with *Q. virginiana* and the two appear to have exchanged bi-directional gene flow recently (tests 16 & 22, Table 2), since the divergence of *Q. virginiana* from the other two Florida clade oaks.

### Distinguishing independent introgression events

Reconstructing the history of introgression among lineages does not translate directly from patterns of shared alleles between them, but instead must be placed in a phylogenetic context. A clear example of this can be seen with *Q. fusiformis*, which appears admixed with respect to every other species of live oak save for its sister taxon *Q. brandegeei* (tests 14-17, Table 2). Of its three potential hybridizing partner lineages it seems least likely to have truly hybridized with *Q. sagraeana*, which is allopatric in Cuba, compared to the other two lineages with which it overlaps in Texas or Mexico. By contrasting these lineages as potential donor lineages using partitioned D-statistics we find that the complex patterns of admixture in *Q. fusiformis* can be explained by a small number of introgression events. The shared derived alleles between *Q. sagraeana* and *Q. fusiformis* in Texas are nearly entirely composed of alleles that these two taxa also share with *Q. virginiana* (Fig. 2B), and similarly, the shared derived alleles between *Q. sagraeana* and *Q. fusiformis* in Mexico are composed almost entirely of alleles also shared with *Q. oleoides* (Fig. 2C). Only *Q. virginiana* shares uniquely introgressed alleles with *Q. fusiformis* in Texas, and only *Q. oleoides* shares uniquely introgressed alleles with *Q. fusiformis* in Mexico. From this we can infer that introgression occurred separately into *Q. fusiformis* from these two distinct lineages, but not from their close relative *Q. sagraeana*, since *Q. sagraeana* does not share introgressed alleles with *Q. fusiformis* to the exclusion of either of its close relatives.

### Hidden ancestry and the Cuban oak

That *Q. sagraeana* would share ancestry with both *Q. oleoides* and the Florida clade oaks to the exclusion of *Q. fusiformis* is consistent with our phylogenetic reconstructions. It is therefore not surprising that introgression from any one of these three related lineages would introduce shared ancestral alleles from all three. By a similar logic, we investigated the origins of the Cuban oak by applying the same test one node lower in the phylogeny - at the first split between a putative ancestor of *Q. oleoides* and the Florida clade - to test which of these two putative parental lineages shares more ancestral (non-introgressed) alleles with *Q. sagraeana*. Our intention, therefore, was to detect evidence of a putative most recent common ancestor (MRCA) whose historical signature has become obscured, by finding evidence of their shared ancestry in alleles that are introgressed from one or more of their descendant lineages into another.

We compared two competing hypotheses: (1) *Q. sagraeana* shares a MRCA with *Q. oleoides* from Central America but subsequently exchanged alleles with one or more Florida clade oaks; or (2) Q. sagraeana shares a MRCA with (or within) the Florida clade oaks but subsequently exchanged genes with *Q. oleoides* (Fig. 3). Both scenarios assume that the ancestral lineage established on Cuba through seed and that later introgression occurred infrequently, either through rare long distance dispersal events or wind-dispersed pollen, most likely at times of low sea level when distances between Cuba and the mainland were reduced.

**Figure 3:**
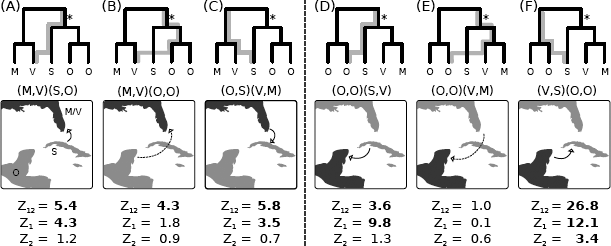
Partitioned D-statistics testing two hypotheses of divergence and gene-flow in the Cuban oak. In hypothesis 1 (A-C) S shares a MRCA with O (light gray on map); in hypothesis 2 (D-F) S shares a MRCA with the Florida clade (taxon abbreviations are as in Fig. 1). An asterisk marks the hypothesized ancestral relationship of S with either lineage. For each scenario sampled tips are shown in the following order (P1,P2)(P3_1_,P3_2_). The direction of introgression being tested is indicated by an arrow on the map, and a gray line traces the path on the topology through which shared ancestral P3 alleles are introduced into P2 to the exclusion of P1. D-statistics are reported as Z-scores.

Partitioning shared versus uniquely derived alleles among these three lineages reveals strong support for the Central American origin hypothesis. If we begin by assuming *Q. oleoides* and *Q. sagraeana* are sister species, we find that *Q. sagraeana* shares a set of uniquely derived alleles with *Q. virginiana* (relative to Q. minima; significant D1), and that a set of derived alleles which putatively arose in the ancestor of *Q. oleoides* and *Q. sagraeana* is also shared with *Q. virginiana* (significant D12), but *Q. oleoides* itself does not share a set of uniquely derived alleles with *Q. virginiana* (non-significant D2) (Fig. 3A; tests 26-31, Table S2). This pattern is consistent with a topology in which *Q. oleoides* and *Q. sagraeana* share a MRCA but introgression occurred from only one descendant lineage. It follows then that if this topology were true all populations of *Q. oleoides* should also share with *Q. virginiana* the set of alleles that arose in the ancestor of *Q. oleoides* and *Q. sagraeana*, despite the fact that *Q. oleoides* never hybridized with *Q. virginiana* directly (they are allopatric). This is precisely what we find (Fig. 3B; tests 32-37, Table S2): shared alleles between Q. oleoides populations are present in *Q. virginiana*, but no single *Q. oleoides* population shows significantly greater genetic similarity with *Q. virginiana*. While this result supports our hypothesized scenario, the true history of divergence and gene flow may be more complex; for example, introgression appears to have also occurred in the reverse direction, from Florida into Cuba, and most likely more than once, since both *Q. virginiana* and *Q. geminata* share a different set of uniquely introgressed alleles with *Q. sagraeana* (Fig. 3C; tests 38-43, Table S2) relative to *Q. oleoides*.

The alternative scenario, in which *Q. sagraeana* is derived from the Florida clade, yields patterns of admixture that are less consistent with the existence of a hypothetical MRCA. This is apparent first in the overabundance of uniquely shared alleles between *Q. sagraeana* and *Q. oleoides* (D_1_), relative to ancestral alleles that should be derived from the hypothetical MRCA of *Q. sagraeana* and *Q. virginiana* (Fig. 3D; tests 44-47, Table S2). It is further apparent because the putative introgression between *Q. sagraeana* and *Q. oleoides* did not introduce any alleles from *Q. virginiana*, or its other Florida clade relatives, which are expected to be introduced alongside alleles from *Q. sagraeana* if they shared a MRCA, and if either acted as an introgressive donor (Fig. 3E; tests 48-53, Table S2). Thus, the strong signal of apparent introgression between *Q. sagraeana* and *Q. oleoides* (Fig. 3F; tests 54-57, Table S2) is most likely, rather, a signal of their shared ancestry made apparent by testing for introgression on an incorrect species tree.

### Demographic models

We further compared these two hypotheses with a third model in which the Cuban population was formed by instantaneous admixture from two parent lineages but remained completely isolated thereafter (Fig. 4A) – a scenario akin to hybrid speciation. By fitting the SFS for these three lineages to demographic models in ∂a∂i (Gutenkunst *et al*. 2009), we found greatest support for a Central American origin (LL=-541.9), followed by the Florida origin (LL=-543.1) and hybrid origin (LL=-555.3) models. The least parameter rich model (hybrid origin) is easily rejected in favor of the two more complex models: the difference in log-likelihood (δ) between models was greater in our observed data than in all simulated data sets generated under the hybrid origin scenario (Fig. 4B). This test was also very sensitive: at a false positive rate of 5%, we had >99% power to reject the hybrid origin model. There is no clear null when comparing the remaining two models to each other, as they are non-nested, and equal in number of parameters. Thus a P-value of 5% may be considered overly stringent (Boettiger *et al*. 2012). The observed *δ* supporting a Central American origin is greater than 93% of simulations generated under the Florida origin model (P=0.07), and using this as our test statistic, we have 92% power to reject a Florida origin if the other model were true. Or, if we use the traditional cutoff of 5%, we have 85% power to correctly distinguish the models (Fig. 4B). Using 2.5x10^−9^ as the average mutation rate per site per generation (inferred from *Populus* (Tuskan *et al*. 2006)), and an average generation time of 30 years, our best model (Central American origin) infers a crown age for these three lineages of 1.75 (1.19-4.00) Mya, with divergence of *Q. sagraeana* occurring 0.19 (0.04-0.31) Mya (Table 3). Introgression occurred predominately into *Q. sagraeana* from the Florida clade, and to a lesser extent from *Q. oleoides*.

**Table 3:** Maximum likelihood (ML) parameter estimates and 95% confidence intervals (CI) for three demographic models for the origin of the Cuban oak.

| Parameter | Model 1 (Florida origin) |  | Model 2 (CA origin) |  | Model 3 (Hybrid origin) |  |
| --- | --- | --- | --- | --- | --- | --- |
|  | ML | 95% CI | ML | 95% CI | ML | 95% CI |
| $N_{MGV}$ ($\times 10^3$) | 89.04 | 71.48–100.27 | 88.34 | 69.72–100.02 | 90.89 | 70.39–104.80 |
| $N_O$ ($\times 10^3$) | 24.52 | 18.59–29.47 | 24.19 | 17.82–29.34 | 28.89 | 22.63–33.64 |
| $N_S$ ($\times 10^3$) | 2.73 | 0.00–5.30 | 8.44 | 2.38–13.46 | 5.76 | 0.34–10.70 |
| $T_{12}$ (Mya) | 1.83 | 1.43–4.00 | 1.75 | 1.19–4.00 | 1.46 | 0.81–3.54 |
| $T_1$ (Mya) | 0.32 | 0.00–0.90 | 0.19 | 0.04–0.31 | 0.06 | 0.00–0.11 |
| $m_{MGV-S}$ ($\times 10^3$) | 0.00 | 0.00–0.01 | 0.00 | 0.00–0.00 | — | — |
| $m_{S-MGV}$ ($\times 10^3$) | 0.18 | 0.02–0.34 | 0.08 | 0.01–0.09 | — | — |
| $m_{S-O}$ ($\times 10^3$) | 0.02 | 0.00–0.03 | 0.06 | 0.02–0.09 | — | — |
| $m_{O-S}$ ($\times 10^3$) | 0.30 | 0.02–0.52 | 0.00 | 0.00–0.00 | — | — |
| $f_{MGV}$ | — | — | — | — | 0.38 | 0.34–0.42 |

## Discussion

Introgressive hybridization is commonly studied at the scale of individual species pairs (Petit *et al.* 1997), among multiple sympatric species (Whittemore & Schaal 1991), or in a sampling of close relatives (The Heliconius Genome Consortium 2012, Gugger & Cavender-Bares 2013, Kane *et al.* 2009, Nadeau *et al.* 2013), but rarely in the context of all extant species within an ecologically and evolutionarily distinct clade. Here, by sampling all relevant populations and comparing them in a phylogenetic context we were able to reconstruct a clade-level history of introgression, and to correct many potentially misleading signals of admixture. We find that every pair of species occurring in close geographic proximity has exchanged some amount of gene flow, with no evidence of introgression that is not concordant with species present day geographic distributions. This suggests that geographic ranges of the live oaks, at least relative to each other, have likely remained stable through time. Such stasis is consistent with the fact that live oak species exhibit substantial differences in adaptations to climatic niche, particularly with regard to drought and freezing tolerances (Cavender-Bares *et al*. 2011, Cavender-Bares & Pahlich 2009, Koehler *et al*. 2012, Ramirez-Valiente et al. In Press, Cavender-Bares *et al*. In Press). Together they span a nearly continuous range from temperate, to dry desert, and even tropical climates. A classic hypothesis for limits on the spread of introgressed alleles between species is that such alleles may facilitate adaptations to intermediate environments within hybrid zones, but decrease fitness elsewhere (Barton & Hewitt 1985). In the live oaks, genetic exchange is theoretically possible throughout a ring-like complex composing up to six interconnected, interfertile species that effectively encircle the Gulf of Mexico, including a connection through Cuba. However, introgressed alleles appear to remain largely concentrated in hybrid zones.

### The comparative nature of tests for introgression

Our analyses demonstrate the difficulty of inferring historical introgression over deep evolutionary time scales. In particular, that sparse sampling can lead to false inferences of hybridization when the source of introgressed alleles is unknown, or stems from multiple sources, as is common for oaks. This is the case for *Q. fusiformis*, which has experienced introgression with two divergent lineages in opposite ends of its geographic range. Because the two lineages with which it hybridized share a common ancestor since their divergence from *Q. fusiformis* each introduced many of the same alleles into it. They also introduced alleles that they share with their other close relatives, including *Q. sagraeana*. Had we failed to sample all extant species, and thus been unable to contrast their patterns of shared versus uniquely derived alleles, we could have easily been misled as to the source of introgression. For example, consider if *Q. oleoides* had not been sampled, in which case only *Q. sagraeana* would appear to share uniquely introgressed alleles with *Q. fusiformis* in the southern part of its range (Fig. 2C); and similarly, a failure to sample the Florida oak clade would lead us to infer introgression from *Q. sagraeana* into *Q. fusiformis* in the northern part of its range (Fig 2B). Given that the true result in each of these cases was that introgression occurred from the most geographically proximate taxon such a distinction may seem trivial. However, if we consider that many studies of introgression focus on only a single species pair, the potential for error, especially in highly diverse clades, is clear. The ability to accurately reconstruct a history of hybridization among multiple closely related species from genomic data would provide an invaluable tool for the study of speciation and reproductive isolation (Rabosky & Matute 2013). The case of the American live oaks makes clear that such histories can be highly complex, and teasing them apart requires both fine-scale sampling and careful hypothesis testing.

### Inferring admixture

We explored a range of methods for detecting introgression and admixture, all of which returned complementary results. *Structure* and *TreeMix* share similarities in their underlying parametric models that infer admixture from the distribution of allele frequencies among populations (Pritchard *et al*. 2000); in the latter case, modeling changes along the branches of a phylogeny (or network) according to genetic drift (Pickrell & Pritchard 2012). The *TreeMix* approach is advantageous over D-statistics in that it takes into account the full phylogeny when inferring admixture, as opposed to individual four or five-taxon subsets of the tree. It thus identifies introgression in the context of all competing hypotheses, and takes into account the non-independence of introgression events. However, when applied to deeply divergent lineages, as in our data, several assumptions of the model may be violated, such as equal population sizes, and that allelic variation arises from ancestral polymorphisms rather than *de novo* mutations (Pickrell & Pritchard 2012). When allowing more than two admixture edges in the live oaks, *TreeMix* inferred one or more instances of introgression between *Q. minima* and the outgroup “population” (tested as various combinations of the four non-*Virentes* white oak taxa), which we suspect is a false result: it is not supported by D-statistics using red oaks as a more distant outgroup [range Z=(0.25–1.99)]. The simplified assumptions underlying non-parametric D-statistics may better facilitate their application for hypothesis testing over deeper evolutionary time scales, however, care must be taken in interpreting results within the context of unsampled phylogenetic relationships.

### Hybrid species

We have focused on reconstructing phylogeny as a representation of the divergence of species through time, assuming that species have remained cohesive lineages despite instances of introgression between them. This view differs from the use of a graph or network to represent truly reticulate histories, or similarly, describing admixed lineages as having arisen through hybrid speciation (Schumer *et al*. 2014). For the latter case, we explicitly tested a model of instantaneous hybrid speciation for the origin of *Q. sagraeana*, the most admixed lineage in the American live oaks. This model was a poor fit compared to one in which an ancestral population of *Q. oleoides* colonized the island and received persistent low levels of introgression from one or more oak species in Florida. A similar scenario in which an island population has undergone nuclear “conversion” towards the genomic makeup of another species has been described for ABC Island brown bears off the coast of Alaska (Cahill *et al*. 2013). Numerous examples of nuclear-chloroplast discordance in mainland oak species suggest this may be a common phenomenon (Petit *et al*. 2004), perhaps exacerbated by limited seed dispersal but widespread pollen flow in oaks.

### Introgression and phylogeny

The effects of introgression on phylogenetic inference are often difficult to detect, but is made easier when multiple individuals are sampled from within a species that vary in their proportions of admixed ancestry. The rare and isolated taxon *Q. brandegeei*, from Baja California, provides an interesting example. Phylogenetic analyses suggested that it is nested within *Q. fusiformis*, appearing more closely related to populations from Mexico than from Texas. This finding, it turns out, is not a result of increased similarity between *Q. brandegeei* and *Q. fusiformis* (Mexico), but rather from the decreased relatedness between *Q. brandegeei* and *Q. fusiformis* (Texas); the latter arising from introgression that occurred into *Q. fusiformis* (Texas) from a more distant clade. This is clear from the phylogenetic results of censored data sets excluding the introgressive donor, which recovered strong support for monophyly of *Q. fusiformis* and its sister relationship to *Q. brandegeei* (Fig. S2E). Should we interpret this to mean that *Q. fusiformis* is not truly paraphyletic with respect to *Q. brandegeei*? The answer depends on what we wish our phylogeny to represent. If it is the historical pattern of population splitting, then *Q. brandegeei* clearly does not belong nested within *Q. fusiformis*. If the phylogeny is meant to show the genetic similarity of sampled individuals, then paraphyly of *Q. fusiformis*, which was recovered in most of our analyses, may be the most appropriate representation.

### The nature of oak species

The nature of species boundaries in oaks is a long-standing topic of philosophical debate. Burger (1975) and later Van Valen (1976) envisioned oaks as a form of “ecological species” in which populations filling a unique ecological niche remain recognizably distinct through shared adaptations regardless of their genomic makeup. Their classic example involves the widespread and easily recognizable bur oak (*Q. macrocarpa*), which hybridizes with up to seven other species across its range. Van Valen conjectured that it does not matter whether a bur oak population in Quebec is more likely to exchange genes with its local congener than with another bur oak population in Texas. He argued that if a recognizably distinct ecological unit persists across this range, it is sufficient to define the species. In the context of more recent views on ecological speciation (Nosil 2012), and the porous nature of species boundaries (Harrison & Larson 2014), the “ecological species” remains relevant, but with an elevated role for genetics - albeit sometimes very few genes (Wu 2001). Our analyses suggest that despite the near continuous geographic distribution of the live oaks, and extensive introgression, species tend to form distinct ecological units that have been maintained over evolutionary time scales.

## Acknowledgments

We thank R. Ree, Y. Brandvain, and the Donoghue Lab at Yale for helpful discussions. This work was partially supported by National Science Foundation grants IOS-0843665 (to J.C-B), DEB-1146380 (to J.C-B & A.R-G), DEB-1146488 (to A.H), and a Lester Armour Graduate Student Fellowship at the Field Museum (to D.E).

---

**Table S1:**
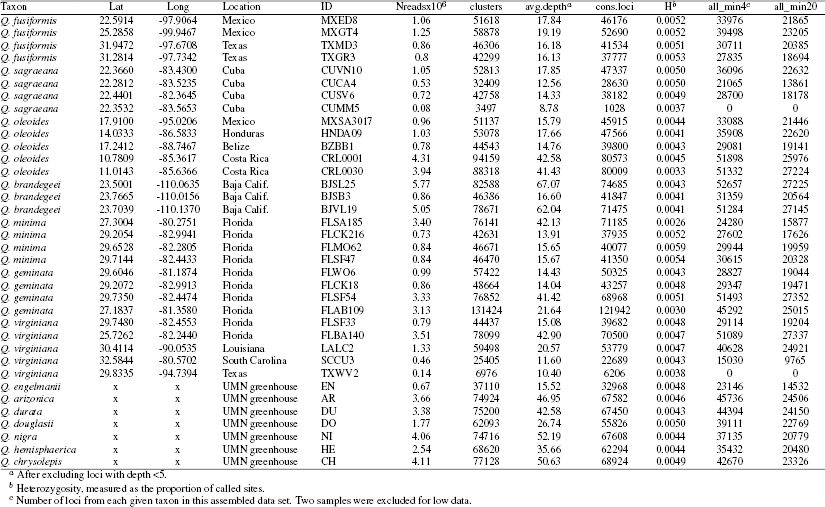
Taxon sampling and summary of RADseq data assembly.

**Table S2:** Selected results of partitioned D-statistic tests investigating the origin of the Cuban oak. Taxon abbreviations are as labeled in Fig. 1, and are arranged such that the dominant signal, when present, is introgression of shared P3 alleles (D_12_) into P2 (ABBBA>BABBA). For each test the corresponding hypothetical scenario from Fig. 3 is indicated. Subscripts show sampling locations for the sampled individual used in each test: *HN* =Honduras, *LA*=Louisiana, *FL*=Florida, *MX*=Mexico.

| test | P1 | P2 | P3<sub>1</sub> | P3<sub>2</sub> | $D_{12}$ | $D_1$ | $D_2$ | $Z_{12}$ | $Z_1$ | $Z_2$ | scenario |
| --- | --- | --- | --- | --- | --- | --- | --- | --- | --- | --- | --- |
| 26 | M | G | S | $O_{HN}$ | 0.10 | 0.14 | 0.00 | 2.9 | 2.5 | 0.0 | A |
| 27 | M | $V_{LA}$ | S | $O_{HN}$ | 0.16 | 0.12 | 0.04 | **5.0** | 2.3 | 0.7 | A |
| 28 | M | $V_{FL}$ | S | $O_{HN}$ | 0.17 | 0.21 | 0.07 | **5.4** | **4.3** | 1.2 | A |
| 29 | G | $V_{LA}$ | S | $O_{HN}$ | 0.08 | 0.01 | 0.07 | 2.8 | 0.2 | 1.3 | A |
| 30 | G | $V_{FL}$ | S | $O_{HN}$ | 0.06 | 0.09 | 0.03 | 1.7 | 1.8 | 0.6 | A |
| 31 | $V_{LA}$ | $V_{FL}$ | S | $O_{HN}$ | 0.01 | 0.09 | -0.02 | 0.2 | 2.0 | 0.3 | A |
| 32 | M | G | $O_{HN}$ | $O_{MX}$ | 0.10 | 0.13 | -0.01 | 2.6 | 1.6 | 0.1 | B |
| 33 | M | $V_{LA}$ | $O_{HN}$ | $O_{MX}$ | 0.14 | 0.06 | 0.09 | **4.1** | 0.8 | 1.4 | B |
| 34 | M | $V_{FL}$ | $O_{HN}$ | $O_{MX}$ | 0.16 | 0.13 | 0.07 | **4.3** | 1.8 | 0.9 | B |
| 35 | G | $V_{LA}$ | $O_{HN}$ | $O_{MX}$ | 0.11 | -0.06 | 0.19 | **3.7** | 1.0 | 2.8 | B |
| 36 | G | $V_{FL}$ | $O_{HN}$ | $O_{MX}$ | 0.07 | -0.05 | 0.11 | 2.1 | 0.7 | 1.7 | B |
| 37 | $V_{LA}$ | $V_{FL}$ | $O_{HN}$ | $O_{MX}$ | -0.03 | 0.01 | -0.05 | 1.2 | 0.1 | 0.8 | B |
| 38 | $O_{HN}$ | S | G | M | 0.22 | 0.23 | 0.11 | **5.6** | **3.4** | 1.8 | C |
| 39 | $O_{HN}$ | S | $V_{LA}$ | M | 0.21 | 0.21 | 0.13 | **6.9** | **3.7** | 2.1 | C |
| 40 | $O_{HN}$ | S | $V_{FL}$ | M | 0.20 | 0.19 | 0.05 | **5.8** | **3.5** | 0.7 | C |
| 41 | $O_{HN}$ | S | $V_{LA}$ | G | 0.25 | 0.20 | 0.25 | **8.0** | **3.8** | **5.0** | C |
| 42 | $O_{HN}$ | S | $V_{FL}$ | G | 0.23 | 0.23 | 0.17 | **7.1** | **4.4** | **3.4** | C |
| 43 | $O_{HN}$ | S | $V_{FL}$ | $V_{LA}$ | 0.25 | 0.21 | 0.12 | **9.8** | **4.1** | 2.2 | C |
| 44 | $O_{MX}$ | $O_{HN}$ | S | G | 0.17 | 0.27 | -0.08 | **5.7** | **7.4** | 1.2 | D |
| 45 | $O_{MX}$ | $O_{HN}$ | S | M | 0.12 | 0.28 | -0.15 | **3.5** | **7.8** | 2.2 | D |
| 46 | $O_{MX}$ | $O_{HN}$ | S | $V_{FL}$ | 0.12 | 0.27 | -0.12 | **3.6** | **9.8** | 1.3 | D |
| 47 | $O_{MX}$ | $O_{HN}$ | S | $V_{LA}$ | 0.11 | 0.28 | -0.07 | **4.1** | **8.4** | 2.2 | D |
| 48 | $O_{MX}$ | $O_{HN}$ | G | M | -0.01 | 0.08 | -0.06 | 0.3 | 1.0 | 0.8 | E |
| 49 | $O_{MX}$ | $O_{HN}$ | $V_{LA}$ | M | -0.01 | -0.02 | 0.03 | 0.2 | 0.3 | 0.4 | E |
| 50 | $O_{MX}$ | $O_{HN}$ | $V_{FL}$ | M | -0.05 | 0.01 | -0.05 | 1.0 | 0.1 | 0.6 | E |
| 51 | $O_{MX}$ | $O_{HN}$ | $V_{LA}$ | G | -0.00 | -0.19 | 0.05 | 0.1 | **3.3** | 0.8 | E |
| 52 | $O_{MX}$ | $O_{HN}$ | $V_{FL}$ | G | 0.02 | -0.12 | 0.04 | 0.5 | 1.6 | 0.7 | E |
| 53 | $O_{MX}$ | $O_{HN}$ | $V_{FL}$ | $V_{LA}$ | -0.01 | 0.08 | -0.06 | 0.3 | 1.0 | 0.8 | E |
| 54 | G | S | $O_{HN}$ | $O_{MX}$ | 0.55 | 0.49 | 0.17 | **26.6** | **11.1** | **3.1** | F |
| 55 | M | S | $O_{HN}$ | $O_{MX}$ | 0.56 | 0.55 | 0.17 | **25.8** | **15.1** | 2.7 | F |
| 56 | $V_{FL}$ | S | $O_{HN}$ | $O_{MX}$ | 0.53 | 0.46 | 0.13 | **26.8** | **12.1** | **3.4** | F |
| 57 | $V_{LA}$ | S | $O_{HN}$ | $O_{MX}$ | 0.52 | 0.48 | 0.12 | **27.3** | **12.4** | 2.7 | F |

**Figure S1:**
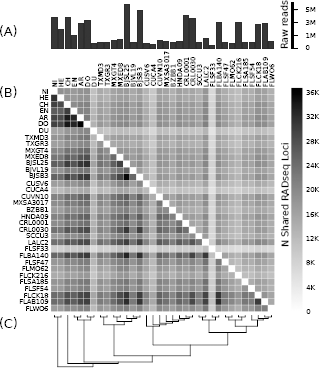
The distribution of shared RADseq loci between samples across two data sets with different thresholds for the minimum sample coverage. (A) The number of raw input reads at the beginning of bioinformatic analyses. (B) Heatmap of locus sharing across the two assembled data sets. The large but sparse “Allmin4” matrix (55.5% missing data) is below the diagonal while the smaller but more complete “Allmin20” matrix (26.6% missing data) is above the diagonal. (C) The inferred “Allmin20” topology.

**Figure S2:**
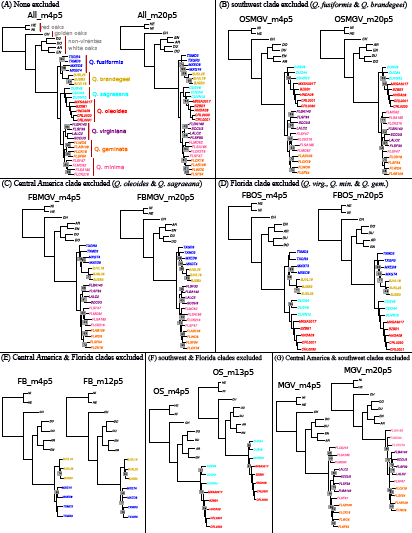
Rooted ML phylogenies inferred from 15 concatenated RADseq data sets. Bootstrap support is 100 except where indicated. Ingroup4t3axon sampling varies among data sets, but each shares the same seven outgroup samples. For each subset of taxa both a sparse and more complete data set were generated. (E-F) Inferred relationships among closely related species or populations are different from the full tree (A) when taxa from distant clades, which may have exchanged genes, are analyzed separately.

**Figure S3:**
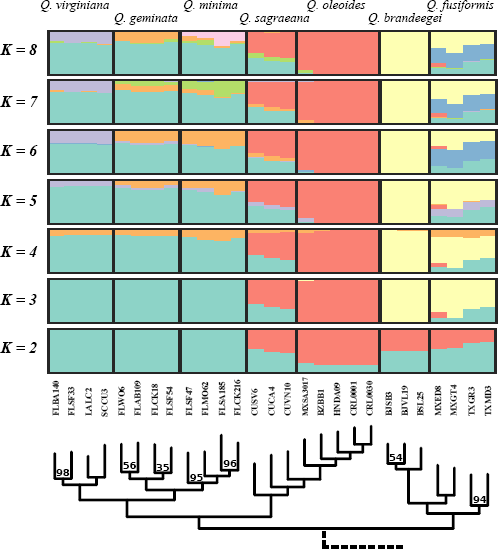
Population clustering with admixture for 27 live oak individuals inferred from 14K SNPs. Specimen IDs are shown. Outgroup taxa were excluded. Clustering was performed at values of *K* between 2–8. The rooted ML tree inferred from the (Allmin4) RADseq data set is also shown for reference. Bootstrap supports are 100 except where indicated.

**Figure S4:**
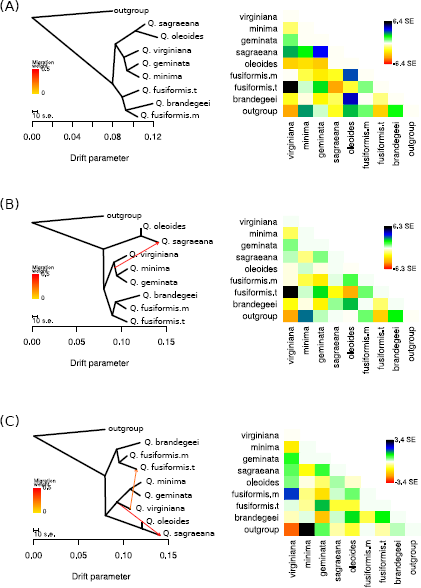
Population splits and admixtures for pooled population samples inferred by TreeMix, and the corresponding allele frequency covariance matrix. (A) A maximum likelihood tree inferred without admixture. (B) The population graph with one admixture edge. (C) The population graph with two admixture edges.

## References

Avise, J.C. (2000). Phylogeography: The History and Formation of Species. Harvard University Press.

Baird, N.A. et al. (2008). Rapid SNP discovery and genetic mapping using sequenced RAD markers. PLoS ONE, 3, e3376.

Barton, N.H. & Hewitt, G.M. (1985). Analysis of hybrid zones. Annual Review of Ecology and Systematics, 16, 113–148.

Boettiger, C., Coop, G. & Ralph, P. (2012). Is your phylogeny informative? measuring the power of comparative methods. Evolution, 66, 2240–2251.

Borgardt, S.J. & Pigg, K.B. (1999). Anatomical and developmental study of petrified *Quercus* (Fagaceae) fruits from the middle Miocene, Yakima Canyon, Washington, USA. American Journal of Botany, 86, 307–325.

Burger, W.C. (1975). The species concept in *Quercus*. Taxon, 24, 45–50.

Cahill, J.A. et al. (2013). Genomic evidence for island population conversion resolves conflicting theories of polar bear evolution. PLoS Genetics, 9, e1003345.

Cavender-Bares, J., Gonzalez-Rodriguez, A., Eaton, D., Hipp, A., Beulke, A. & Manos, P. (In Press). Phylogeny and biogeography of the American live oaks *(Quercus* subsection *Virentes*): A genomic and population genetic approach. Molecular Ecology.

Cavender-Bares, J., Gonzalez-Rodriguez, A., Pahlich, A., Koehler, K. & Deacon, N. (2011). Phylogeography and climatic niche evolution in live oaks *(Quercus* series *Virentes)* from the tropics to the temperate zone. Journal of Biogeography, 38, 962–981.

Cavender-Bares, J. & Pahlich, A. (2009). Molecular, morphological, and ecological niche differentiation of sympatric sister oak species, *Quercus virginiana* and *Q. geminata* (Fagaceae). American Journal of Botany, 96, 1690–1702.

Coyne, J.A. & Orr, H.A. (2004). Speciation. W.H. Freeman.

Dumolin-Lapegue, S., Kremer, A. & Petit, R.J. (1999). Are chloroplast and mitochondrial DNA variation species independent in oaks? Evolution, 53, 1406–1413.

Durand, E.Y., Patterson, N., Reich, D. & Slatkin, M. (2011). Testing for ancient admixture between closely related populations. Molecular Biology and Evolution, 28, 2239–2252.

Earl, D.A. & vonHoldt, B.M. (2012). STRUCTURE HARVESTER: a website and program for visualizing STRUCTURE output and implementing the Evanno method. Conservation Genetics Resources, 4, 359–361.

Eaton, D.A.R. (2014). PyRAD: assembly of de novo RADseq loci for phylogenetic analyses. Bioinformatics, 30, 1844–1849.

Eaton, D.A.R. & Ree, R.H. (2013). Inferring phylogeny and introgression using RADseq data: An example from flowering plants (Pedicularis: Orobanchaceae). Systematic Biology, 62, 689–706.

Green, R.E., Krause, J., Briggs, A.W., Maricic, T., Stenzel, U., Kircher, M., Patterson, N., Li, H., Zhai, W., Fritz, M.H.Y., Hansen, N.F., Durand, E.Y., Malaspinas, A.S., Jensen, J.D., Marques-Bonet, T., Alkan, C., Prüfer, K., Meyer, M., Burbano, H.A., Good, J.M., Schultz, R., Aximu-Petri, A., Butthof, A., Höber, B., Höffner, B., Siegemund, M., Weihmann, A., Nusbaum, C., Lander, E.S., Russ, C., Novod, N., Affourtit, J., Egholm, M., Verna, C., Rudan, P., Brajkovic, D., Kucan, v., Gu, I., Doronichev, V.B., Golovanova, L.V., Lalueza-Fox, C., Rasilla, M.d.l., Fortea, J., Rosas, A., Schmitz, R.W., Johnson, P.L.F., Eichler, E.E., Falush, D., Birney, E., Mullikin, J.C., Slatkin, M., Nielsen, R., Kelso, J., Lachmann, M., Reich, D. & Pääbo, S. (2010). A draft sequence of the Neandertal genome. Science, 328, 710–722.

Gugger, P.F. & Cavender-Bares, J. (2013). Molecular and morphological support for a Florida origin of the Cuban oak. Journal of Biogeography, 40, 632–645.

Gutenkunst, R.N., Hernandez, R.D., Williamson, S.H. & Bustamante, C.D. (2009). Inferring the joint demographic history of multiple populations from multidimensional SNP frequency data. PLoS Genetics, 5, e1000695.

Hardin, J.W. (1975). Hybridization and introgression in *Quercus alba*. Journal of the Arnold Arboretum, 56, 336–363.

Harrison, R.G. & Larson, E.L. (2014). Hybridization, introgression, and the nature of species boundaries. Journal of Heredity, 105, 795–809.

Hipp, A.L., Eaton, D.A.R., Cavender-Bares, J., Fitzek, E., Nipper, R. & Manos, P.S. (2014). A framework phylogeny of the American oak clade based on sequenced RAD data. PLoS ONE, 9, e93975.

Hudson, R.R. (2002). Generating samples under a Wright-Fisher neutral model of genetic variation. Bioinformatics, 18, 337–338.

Jakobsson, M. & Rosenberg, N.A. (2007). CLUMPP: a cluster matching and permutation program for dealing with label switching and multimodality in analysis of population structure. Bioinformatics, 23, 1801–1806.

Kane, N.C., King, M.G., Barker, M.S., Raduski, A., Karrenberg, S., Yatabe, Y., Knapp, S.J. & Rieseberg, L.H. (2009). Comparative genomic and population genetic analyses indicate highly porous genomes and high levels of gene flow between divergent *Helianthus* species. Evolution, 63, 2061–2075.

Koehler, K., Center, A. & Cavender-Bares, J. (2012). Evidence for a freezing tolerance-growth rate trade-off in the live oaks *(Quercus* series *Virentes)* across the tropical-temperate divide. The New Phytologist, 193, 730–744.

Kurz, H. & Godfrey, R.K. (1962). Trees of Northern Florida. University of Florida Press.

Leaché, A.D., Harris, R.B., Rannala, B. & Yang, Z. (2014). The influence of gene flow on species tree estimation: a simulation study. Systematic Biology, 63, 17–30.

Maddison, W.P. & Knowles, L.L. (2006). Inferring phylogeny despite incomplete lineage sorting. Systematic Biology, 55, 21–30.

Muller, C.H. (1961). The live oaks of the series *Virentes*. American Midland Naturalist, 65, 17–39.

Nadeau, N.J. et al. (2013). Genome-wide patterns of divergence and gene flow across a butterfly radiation. Molecular Ecology, 22, 814–826.

Nixon, K. & Muller, C. (1997). *Quercus* Linnaeus sect. *Quercus* white oaks. In: Flora of North America North of Mexico. Oxford University Press, New York, pp. 436–506.

Nixon, K.C. (1984). A biosystematic study of Quercus series Virentes (the live oaks) with phylogenetic analyses of Fagales, Fagaceae and Quercus. Ph.D. thesis, University of Texas.

Nosil, P. (2012). Ecological Speciation. Oxford University Press, Oxford; New York.

Pearse, I.S. & Hipp, A.L. (2009). Phylogenetic and trait similarity to a native species predict herbivory on non-native oaks. Proceedings of the National Academy of Sciences, 106, 18097–18102.

Pérez, F. & Granger, B.E. (2007). IPython: a system for interactive scientific computing. Computing in Science and Engineering, 9, 21–29.

Petit, R.J., Bodénès, C., Ducousso, A., Roussel, G. & Kremer, A. (2004). Hybridization as a mechanism of invasion in oaks. New Phytologist, 161, 151–164.

Petit, R.J. & Excoffier, L. (2009). Gene flow and species delimitation. Trends in Ecology & Evolution, 24, 386–393.

Petit, R.J. et al. (1997). Chloroplast DNA footprints of postglacial recolonization by oaks. Proceedings of the National Academy of Sciences, 94, 9996–10001.

Pickrell, J.K. & Pritchard, J.K. (2012). Inference of population splits and mixtures from genome-wide allele frequency data. PLoS Genetics, 8, e1002967.

Pritchard, J.K., Stephens, M. & Donnelly, P. (2000). Inference of population structure using multilocus genotype data. Genetics, 155, 945–959.

Rabosky, D.L. & Matute, D.R. (2013). Macroevolutionary speciation rates are decoupled from the evolution of intrinsic reproductive isolation in Drosophila and birds. Proceedings of the National Academy of Sciences, 110, 15354–15359.

Ramirez-Valiente, J., Koehler, K. & Cavender-Bares, J. (In Press). Inter- and intraspecific variation in xanthophyll cycle pigments and anthocyanin accumulation in response to drought and low temperature in live oaks *(Quercus* series *Virentes*). New Phytologist.

Rhymer, J.M. & Simberloff, D. (1996). Extinction by hybridization and introgression. Annual Review of Ecology and Systematics, 27, 83–109.

Rogers, A.R. & Bohlender, R.J. (In Press). Bias in estimators of archaic admixture. Theoretical Population Biology.

Schumer, M., Rosenthal, G.G. & Andolfatto, P. (2014). How common is homoploid hybrid speciation? Evolution, 68, 1553–1560.

Stamatakis, A. (2014). RAxML version 8: a tool for phylogenetic analysis and post-analysis of large phylogenies. Bioinformatics, 30, 1312–1313.

The Heliconius Genome Consortium (2012). Butterfly genome reveals promiscuous exchange of mimicry adaptations among species. Nature, 487, 94–98.

Tuskan, G.A. et al. (2006). The genome of black cottonwood, *Populus trichocarpa* (Torr. & Gray). Science, 313, 1596–1604.

Van Valen, L. (1976). Ecological species, multispecies, and oaks. Taxon, 25, 233–239.

Whittemore, A.T. & Schaal, B.A. (1991). Interspecific gene flow in sympatric oaks. Proceedings of the National Academy of Sciences of the United States of America, 88, 2540–2544.

Wu, C.I. (2001). The genic view of the process of speciation. Journal of Evolutionary Biology, 14, 851–865.

